# A Comparative Study of Epithelial Transport in the Rat and Mouse Kidneys: Modelling and Analysis

**DOI:** 10.1101/2022.07.12.499702

**Authors:** Anita T. Layton

**Affiliations:** University of Waterloo

## Abstract

The goal of the present study was to investigate the functional implications of sex and species differences in the pattern of transporters along nephrons in the rat and mouse kidney, as reported by Veiras et al. (*J Am Soc Nephrol* 28: 3504–3517, 2017). To do so, we developed the first sex-specific computational models of epithelial water and solute transport along the nephrons from male and female mouse kidneys, and conducted simulations along with our published rat models. These models account for the sex differences in the abundance of apical and basolateral transporters, glomerular filtration rate, and tubular dimensions. Model simulations predict that 73% and 57% of filtered Na^+^ is reabsorbed by the proximal tubules of male and female rat kidneys, respectively. Due to their smaller transport area and lower NHE3 activity, the proximal tubules in the mouse kidney reabsorb a significantly smaller fraction of the filtered Na^+^, at 53% in male and only 34% in female. The lower proximal fractional Na^+^ reabsorption in female kidneys of both rat and mouse is due primarily to their smaller transport area, lower Na^+^/H^+^ exchanger activity, and lower claudin-2 abundance, culminating in significantly larger fractional delivery of water and Na^+^ to the downstream nephron segments in female kidneys. Conversely, the female distal nephron exhibits a higher abundance of key Na^+^ transporters, including Na^+^-Cl^−^ cotransporters in both species, epithelial Na^+^ channels for the female rat, and Na^+^-K^+^-Cl^−^cotransporters for the female mouse. The higher abundance of transporters accounts for the enhanced water and Na^+^ transport along the female rat and mouse distal nephrons, relative to the respective male, resulting in similar urine excretion between the sexes. Model simulations indicate that the sex and species differences in renal transporter patterns may partially explain the experimental observation that, in response to a saline load, the diuretic and natriuretic responses were more rapid in female rats than males, but no significant sex difference was found in mice. These computational models can serve as a valuable tool for analyzing findings from experimental studies conducted in rats and mice, especially those involving genetic modifications.

## Introduction

Almost half of the adult population in the United States is now classified as hypertensive, according to the guidelines that define hypertension as a systolic blood pressure above 130 mmHg and a diastolic blood pressure greater than 80 mmHg. Arterial blood pressure is determined by extracellular fluid volume and total body Na^+^ (21). The balance of fluid and electrolytes is regulated by the kidneys via its transport mechanisms that mediate Na^+^ reabsorption. As such, the kidneys play a crucial role in long-term blood pressure control. Renal transporters that play a crucial role in reabsorbing the approximately 1.5 kg of salt are reabsorbed daily in humans include, (i) along the proximal tubules, the apical Na^+^/H^+^ exchanger 3 (NHE3), (ii) along the thick ascending limb of the loop of Henle, the apical Na^+^-2Cl^-^-K^+^ cotransport (NKCC2), and (iii) along the distal convoluted tubules, the apical Na^+^-Cl^−^ cotransporter (NCC). Na^+^ transport is driven by the basolateral Na^+^-K^+^-ATPase. The final fine-tuning of Na^+^ transport is mediated by the collecting duct principal cell epithelial Na^+^ channel (ENaC) (24). Dysregulation of these transporters and channels can result in hypotension or hypertension.

Throughout much of their life, men have higher blood pressure than (31). This sex difference has been reported across race, and across species including rats, mice and chickens (27), dogs (35), as well as animal models of hypertension (28, 33). Hence, there is much interest in gaining a better understanding of the mechanisms controlling blood pressure in both sexes. Given the kidney’s key role in blood pressure regulation, sex differences in hypertension may be attributable, in part, to sex differences in kidney structure and function. Indeed, the renal electrolyte transporters and channels that mediate electrolyte transport exhibits abundance patterns that are sexually dimorphic, as reported by Veiras et al. (36). Their findings revealed markedly different transport capacity in the proximal tubule of the male and female nephrons of the rat and mouse. Notably, female rat proximal tubules exhibit greater phosphorylated (inactive) NHE3 and re-distribution to the base of the microvilli where activity is lower (3), whereas female mice exhibit a lower NHE3 abundance. In both cases, NHE3 activity is lower in females. Consequently, relative to males, in females the proximal tubule reabsorbs a substantially lower fraction of filtered Na^+^ (36). The higher fractional Na^+^ distal delivery in females is handled by the augmented transport capacity in the downstream segments. Despite these overall similarities, there are interesting inter-species differences. Proximal tubule apical water permeability is lower in female rat compared to male rat, but higher in female mouse compared to male mouse. While both female rat and mouse exhibit higher abundance of NCC and claudin-7 distal along the distal nephron segments exhibit, female rats also exhibit more abundant ENaC, whereas female mice express more NKCC (36).

What are the functional implications of the sexual dimorphism in renal transporter and channel expression patterns in rodents? What are the functional implications of the inter-species differences? To answer the first question for the rat, we have previously developed sex-specific computational models of solute and water transport along the rat nephrons (9, 10, 19), and applied those models to investigate sex differences in kidney function and in responses to pharmacological manipulations (12, 14, 29). In this study, we developed the first sex-specific computation models for the nephron transport in the mouse kidneys. Using these models, we conduct a comparative study of kidney function in rats and mice. The mouse models in particular can also serve as a valuable tool in better understanding findings in knock-out studies.

## Methods

We have previously published epithelial cell-based models of solute and water transport along the nephrons in a rat kidney (13, 16). Models were formulated specifically for the male rat and female rat, to capture the sex differences in morphology, hemodynamics, and transporter pattern. In this study, we developed analogous models for the male and female mice. Model equations (see Ref. (43)) are based on mass conservation which are valid for both sexes and across species. Hence, those same equations are used in the mouse models, but appropriate parameter values are changed to account for interspecies and sex differences. The parameter changes for the male and female mouse models versus rats are summarized in Table 1. Each rat kidney model is assumed to contain 36,000 nephrons, whereas each mouse kidney model has 24,000 nephrons.

**Table 1.**
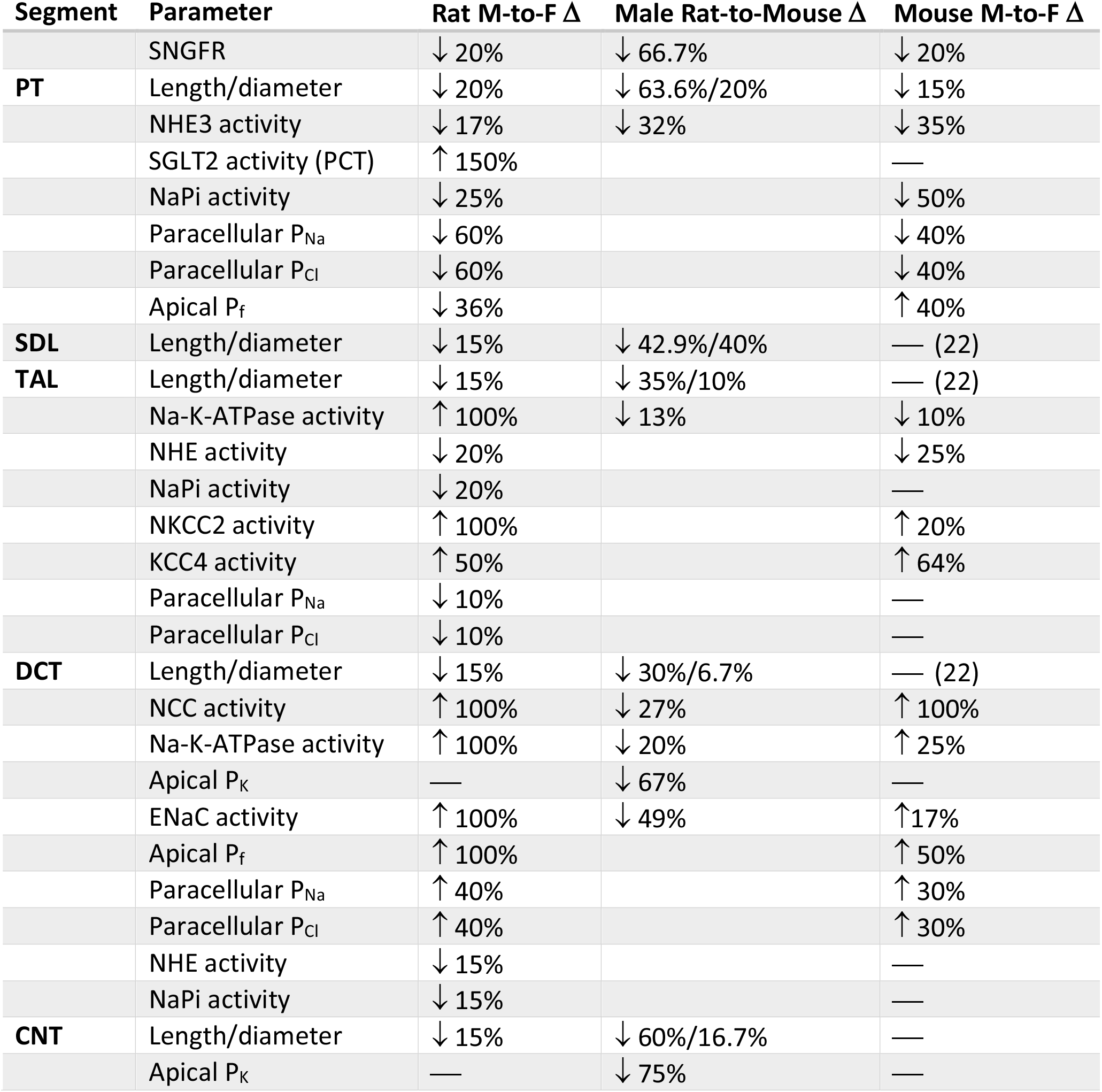
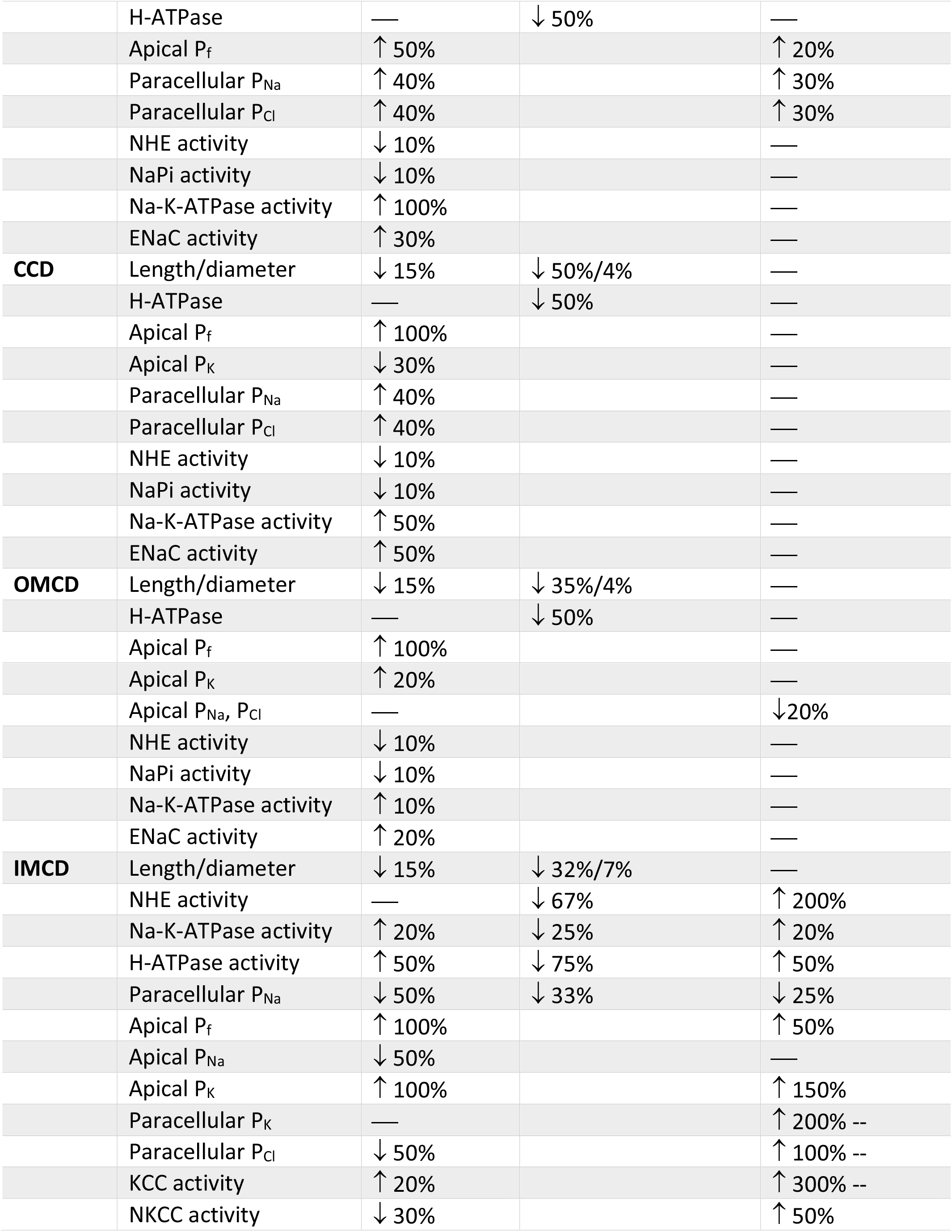
Key parameter differences between male rat and male mouse nephron models, between male and female rat nephron models, and between male and female mouse nephron models. Rat parameters are taken from published models (9, 10). Mouse parameters are based on data from Refs. (22, 32, 36, 40).

### Model structure

Rodent kidneys consist of multiple types of nephrons; superficial and juxtamedullary nephrons make up most of them (80–82). In a rat, about two-thirds of the nephron population are superficial nephrons; the loops of Henle of superficial nephrons turn before reaching the inner medulla. The remaining third are called juxtamedullary nephrons. In a mouse, the ratio of superficial-to-juxtamedullary nephron is estimated to be 82:11 (41). The loops of Henle of juxtamedullary nephrons reach into differing depths of the inner medulla. To capture this heterogeneity, we represent six classes of nephrons: a superficial nephron (denoted by “SF”) and five juxtamedullary nephrons that are assumed to reach depths of 1, 2, 3, 4, and 5 mm (denoted by “JM-1”, “JM-2”, “JM-3”, “JM-4”, and “JM-5”, respectively) into the inner medulla. For the rat, the ratios for the six nephron classes are *n*_*SF*_ = 2/3, *n*_*JM*−1_ = 0.4/3, *n*_*JM*−2_ = 0.3/3, *n*_*JM*−3_ (= 0.15/3, *n*_*JM*−4_) = 0.1/3, and *n*_*JM*−5_ = 0.05/3(11); for the mouse, *n*_*SF*_ = 0.82, *n*_*JM*−1_ = 0.82 × 0.4, *n*_*JM*−2_ = 0.82 × 0.3, *n*_*JM*−3_ = 0.82 × 0.15, *n*_*JM*−4_ = 0.82 × 0.1, and *n*_*JM*−5_ = 0.82 × 0.05(41). The model superficial nephron includes the proximal tubule, short descending limb, thick ascending limb, distal convoluted tubule, and connecting tubule segments. Each of the model juxtamedullary nephron includes all the same segments of the superficial nephron with the addition of the long descending limbs and ascending thin limbs; these are the segments of the loops of Henle that extend into the inner medulla. The length of the long descending limbs and ascending limbs are determined by which type of juxtamedullary nephron is being modeled. The connecting tubules of the five juxtamedullary nephrons and the superficial nephron coalesce into the cortical collecting duct. SNGFR for juxtamedullary nephrons is higher than the superficial nephron SNGFR. In rats, the juxtamedullary SNGFR is taken to be 40-50% higher than the superficial SNGFR (23, 26); in mice, the difference is 20%, based on superficial SNGFR and GFR measurements (22, 40).

Each nephron segment is modeled as a tubule lined by a layer of epithelial cells in which the apical and basolateral transporters vary depending on the cell type (i.e., segment, which part of segment, intercalated and principal cells). The model accounts for the following 15 solutes: Na^+^, K^+^, Cl^-^, HCO_3_^-^, H_2_CO_3_, CO_2_, NH_3_, NH_4_^+^, HPO_4_^2-^, H_2_PO_4_^-^, H^+^, HCO_2_^-^, H_2_CO_2_, urea, and glucose. The model consists of a large system of coupled ordinary differential and algebraic equations, solved for steady state, and predicts luminal fluid flow, hydrostatic pressure, membrane potential, luminal and cytosolic solute concentrations, and transcellular and paracellular fluxes through transporters and channels. A schematic diagram of the various cell types, with interspecies differences highlighted, is given in Fig. 1.

**Figure 1.**
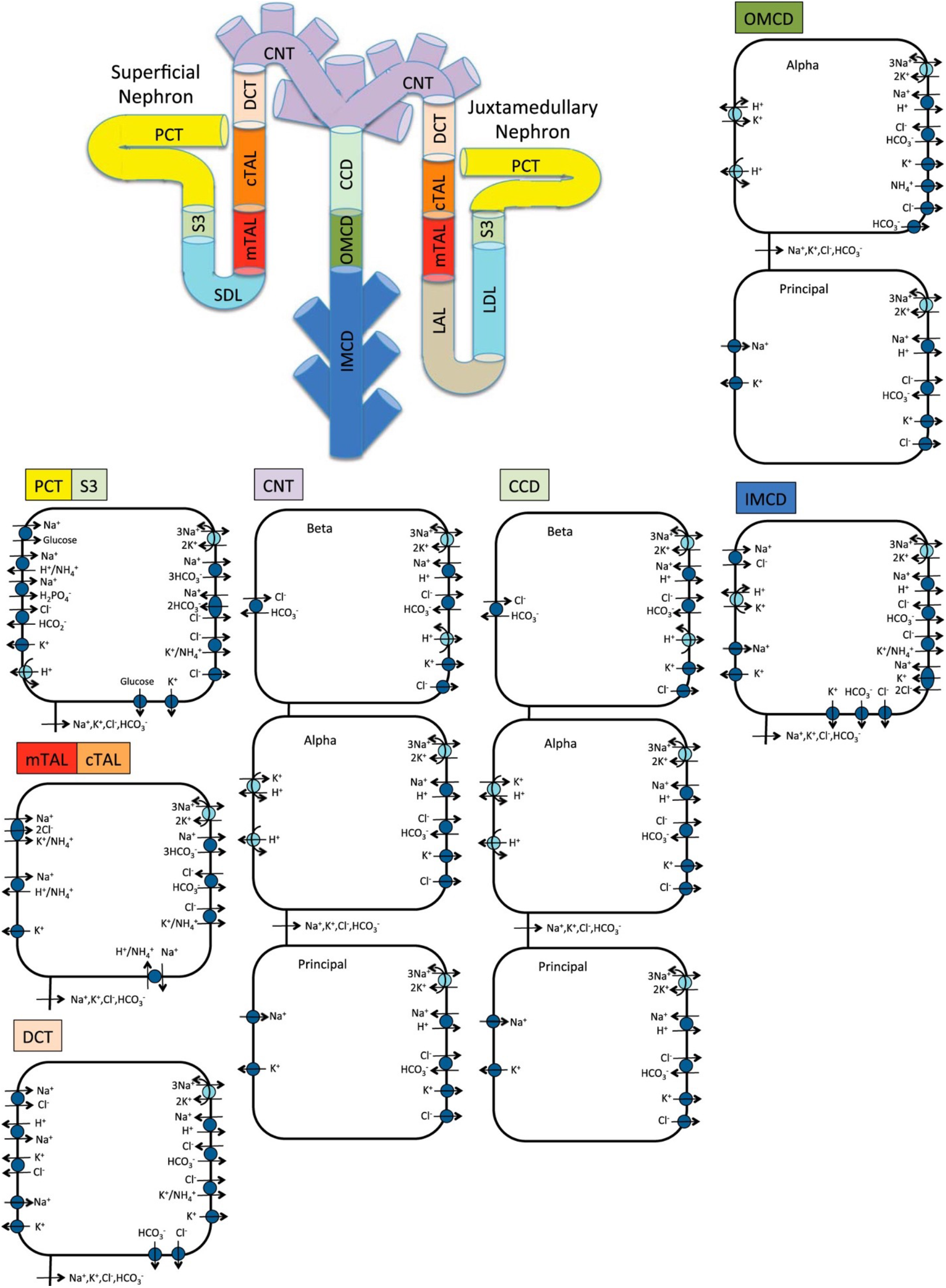
Schematic diagram of the nephron system (not to scale). Model structure is similar for the rat and mouse. The model includes one representative superficial nephron and five representative juxtamedullary nephrons, each scaled by the appropriate population ratio. Only the superficial nephron and one juxtamedullary nephron are shown. Along each nephron, the model accounts for the transport of water and 15 solutes (see text). The distal tubule is divided into the early and late portions of the distal convoluted tubule (DCT). The diagram displays only the main Na^+^, K^+^, and Cl^−^ transporters. PCT, proximal convoluted tubule; SDL, short or outer medullary descending limb; mTAL, medullary thick ascending limb; cTAL, cortical thick ascending limb; CNT, connecting tubule; CCD, cortical collecting duct; OMCD, outer-medullary collecting duct; IMCD, inner medullary collecting duct; LDL, thin descending limb; LAL, thin ascending limb. Reprinted with permission from Ref. (16).

### Model equations

Here we briefly describe the epithelial transport model, which has been described in more detail in Refs. (41,59). A schematic diagram of the epithelial transport model and the segments is given in **Error! Reference source not found**..

Consider a segment denoted *i*, (*i =*PCT, S3, etc.) at steady state. For a specific cell along the nephron the cellular, paracellular, and luminal space are represented by different compartments. Let 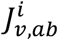 denote the transmembrane volume flux from compartment *a* to compartment *b* calculated using the osmotic and hydraulic pressure by

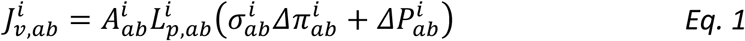

where 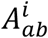 is the membrane area, 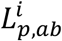 denotes the hydraulic permeability, 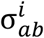 is the reflection coefficient, 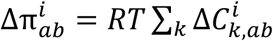, where 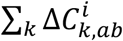 is the osmolality difference, 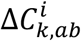 denotes the concentration gradient of solute *k* along the membrane, *R* the ideal gas constant, *T* the thermodynamic temperature, and 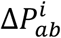 is the hydrostatic pressure gradient.

Conservation of volume in the cellular and paracellular compartments (denoted by *C* and *P* respectively) is given by

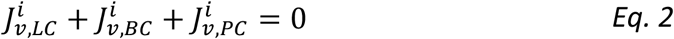

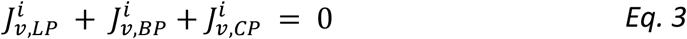

where *L* and *B* denote the lumen and blood, respectively. In the lumen, for non-coalescing segments, conservation of volume is given by

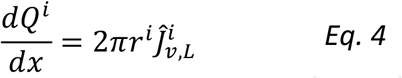

where *Q*^*i*^ denotes the volume flow, *r*^*i*^ is the luminal radius, and 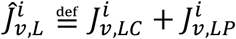 is the total volume flux. The luminal fluid flow is given by pressure-driven Poiseuille flow

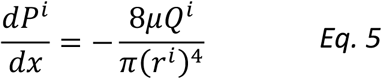

where *P*^*i*^ is the hydrostatic pressure and μ is the luminal fluid viscosity.

The luminal radius *r*^*i*^ is assumed to be rigid for all the segments except the proximal tubule (i.e., *r*^*i*^ is constant). Flow-dependent transport for a compliant tubule is assumed along the proximal tubule using the torque-modulated effects approach developed in in Ref. (60) and described briefly here.

The compliant luminal radius in the proximal tubule (*r*_*PT*_) is given by

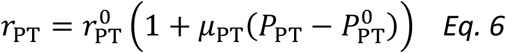

where 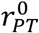 is the reference radius, 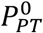 is the reference *pressure, and μ*_*PT*_ characterizes tubular compliance. In the proximal tubule, transporter density is modulated by luminal flow, so that the microvillous torque is given by

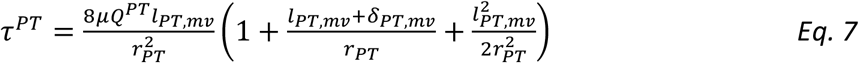

where *l*_*PT,mv*_ is the microvillous length and *δ*_*PT,mv*_ denotes the height above the microvillous tip where drag is considered. Using this, we compute

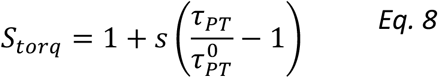

where 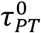 is computed using *Eq. 7* with *Q*^*PT*^ equal to the initial volume flow at the start of the proximal tubule and *r*_PT_ as the initial proximal tubule radius, *s* is a scaling factor. The flux of solutes and volume out of the lumen is then scaled by *S*_*torq*_ to capture the torque-modulated effects.

For coalescing segments, i.e., the CNT and IMCD, water and solute flows are scaled by the tubule population where *ω*^*i*^ (for *i =* CNT or IMCD) denotes the fraction of the tubule remaining at a given spatial location where

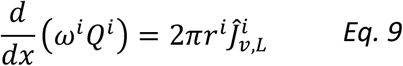

denotes the conservation of volume in the lumen.

The fraction of connecting tubules *ω*^*CNT*^ remaining at coordinate *x*^*CNT*^ (determined as distance from the start of the connecting tubule) is given by

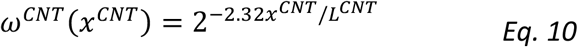

where *L*^*CNT*^ is the connecting tubule length.

The collecting ducts also coalesce in the inner medulla so that the fraction of collecting ducts at distance from the start *x*^*IMCD*^ is given by

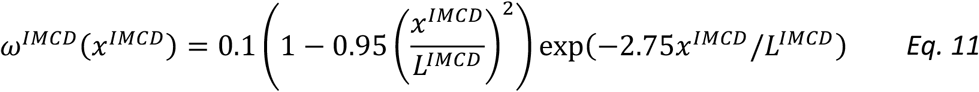

where *L*^*IMCD*^ is the length of the inner-medullary collecting duct. *Eq*. 10 and *Eq. 11* are derived in Ref. (61).

Next, conservation of mass for a non-reacting solutes *k* in the cellular and paracellular space is given by

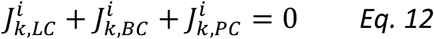

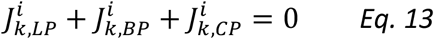

where 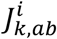 denotes the transmembrane flux of solute *k* from compartment *a* to compartment *b*. Unlike transmembrane volume flux (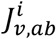, described above), transmembrane solute flux 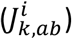 depends on electro diffusion, coupled transport, and primary active transport across ATP-driven pumps. In general, for a solute *k* across membrane *a,b* the solute flux 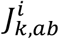 is given by

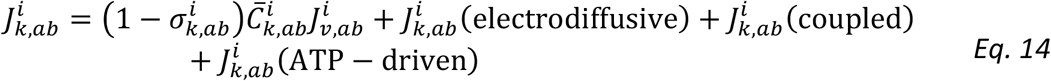

where the first term represents convective transport, 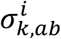 is the reflection coefficient, and

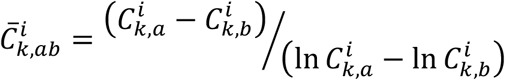

is the mean membrane solute concentration. The second term in *Eq. 14* denotes the transport via electrodiffusive flux. For ions, electrodiffusive flux is given by the Goldman-Hodgkin-Katz equation

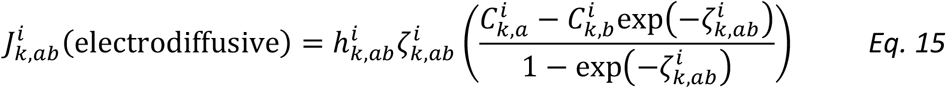

where 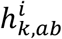 is the membrane permeability for solute *k*,

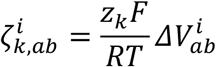

where *z*_*k*_ is the solute valence, *F* is Faraday’s constant, and 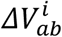 denotes the electrical potential difference. For an uncharged solute, the electrodiffusive flux is given by simple diffusion

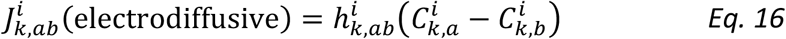

since transport does not depend on the electric potential, i.e., only depends on the concentration gradient.

The third term in *Eq. 14* denotes the total coupled transport of a solute *k* across cotransporters and exchangers. The final term in *Eq. 14* denotes the total transport via primary active transport (i.e., across ATPases). Transport via cotransporters, exchangers, and primary active transport is determined using existing kinetic models for each transporter (59). **Error! Reference source not found**. shows key transporters for the cells of individual segments.

In the lumen, conservation of non-reacting solutes is given by

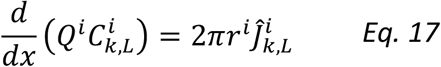

for non-coalescing segments and

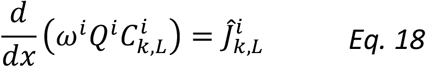

for coalescing segments, where 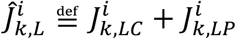 denotes the total flux of solute *k* and *ω*^*i*^ is as defined in *Eq. 10* and *Eq. 11* for the CNT and IMCD respectively.

Conservation is imposed in a similar way on the total buffers for reacting solutes, i.e., acid-base pairs (HPO_4_^2-,^ H_2_PO_4_^-^), (NH_3_, NH_4_^+^) or (HCO_2_^-^, H_2_CO_2_). Conservation for reacting solutes is different because of their net generation from reactions. Details on conservation of reacting solutes can be found in Refs. (59,62).

## Results

We simulated nephron transport function for the male and female rats, and for the male and female mice. Figure 2 shows the predicted total delivery of key solutes [Na^+^, K^+^, Cl^−^, HCO_3_^−^, and NH_4_^+^] and fluid to the inlets of individual nephron segments, expressed per kidney, in the four models. Recall that GFR in lower in the female models; thus, solute delivery into the proximal tubules is also lower in females. Note also that plasma [K^+^] is lower in female rodents.

**Fig. 2.**
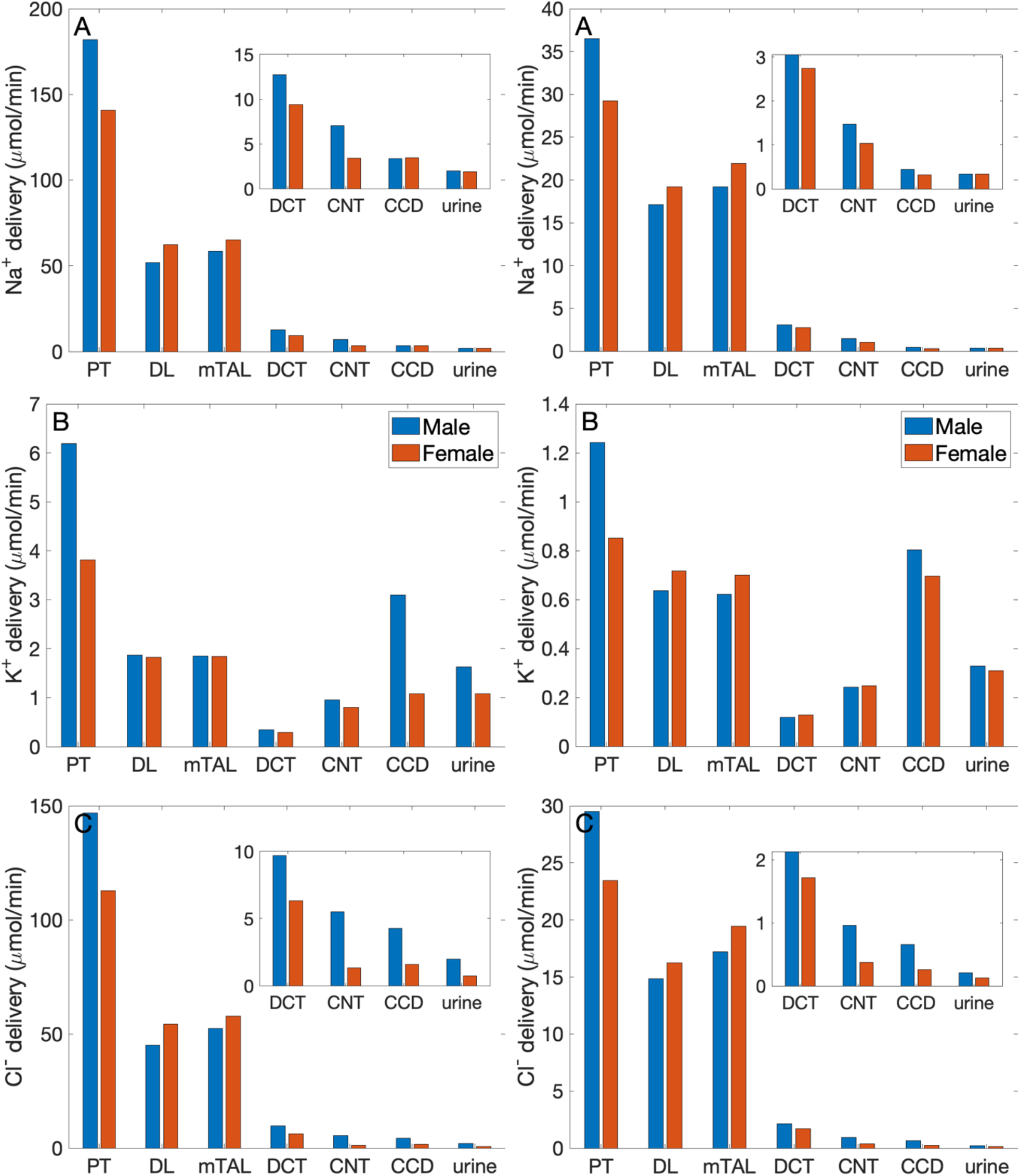
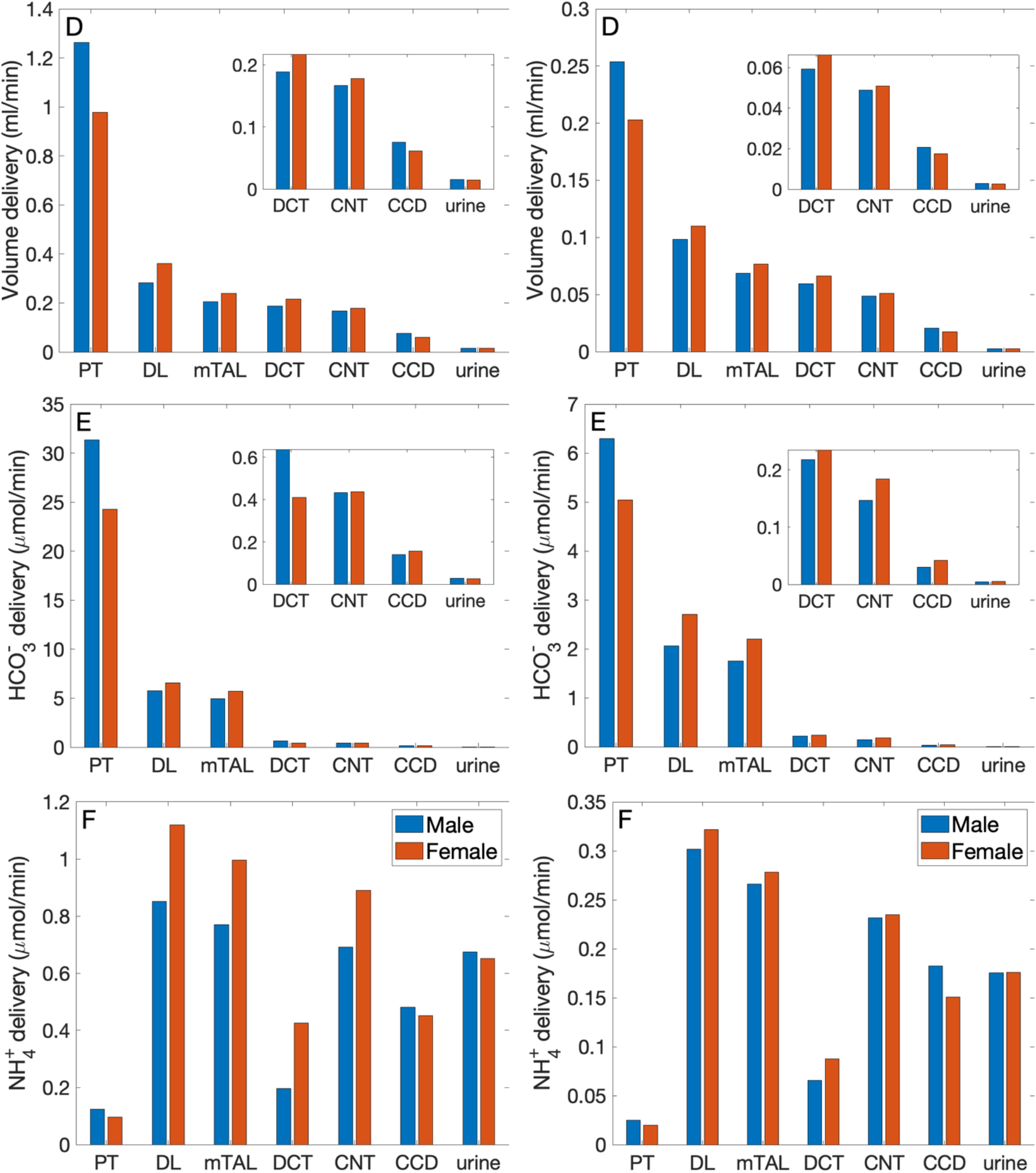
Delivery of key solutes and fluid to the beginning of individual nephron segments in male and female rats (left column) and mice (right columns). Flows along superficial and juxtamedullary are scaled by their population and added together. The rightmost sets of bars show urine excretion rates. PT, proximal tubule; DL, descending limb; mTAL, medullary thick ascending limb; DCT, distal convoluted tubule; CNT, connecting tubule; CCD, cortical collecting duct. Insets: reproductions of distal segment values. The difference between inflow of consecutive segments corresponds to net transport along that segment. Notably proximal tubules mediate significantly more solute and water transport in rats than mice; and within each species, more in males than females.

### Rats versus mice

The predictions of the rat and mouse nephron models share a number of similarities. Almost all the filtered Na^+^, Cl^-^, HCO_3_^-^, and water is reabsorbed. Most of the Na^+^ reabsorption happens along the proximal tubules, via the NHE3, and the thick ascending limbs, via the NKCC2 and Na^+^-K^+^-ATPase. Na^+^ transport is accompanied by Cl^-^ and, along water-permeable segments, by water. The majority of the filtered K^+^ is reabsorbed along the proximal tubules and thick ascending limbs. Downstream of Henle’s loops, the connecting tubules vigorously secrete K^+^. In both rodent models, the proximal tubule is a major site of NH_4_^+^ secretion via substitution of H^+^ in the NHE3 transporter and secretion of NH_3_. A substantial fraction of NH_4_^+^ is then reabsorbed along the thick ascending limbs by substituting for K^+^ in NKCC2.

The most notable difference between solute handling in the rat and mice is arguably the significantly smaller fraction of filtered Na^+^ reabsorbed along the mouse proximal tubules, compared to the rat. This interspecies difference can be attributed to the smaller transport area and lower NHE3 activity in mice. Recall that SNGFR of a superficial mouse nephron is taken to be 1/3 that of a rat (40). If tubular transport area in the mouse models is also scaled down to 1/3, then taken in isolation, the mouse models would predict urinary excretion rates that are about 1/3 of the corresponding rat models, essentially acting like a smaller rat. The total transport area of a given segment is proportional to the product of its length and diameter. Transport area of the mouse PT is approximately one-third that of the sex-matched rat. That, together with the lower NHE3 activity (Table 1), results in the mouse PT transporting a significantly smaller fraction of the filtered Na^+^ than rat in both sexes. The rat proximal tubules are predicted to reabsorb 73% and 57% of the filtered Na^+^ in the male and female models, respectively. In contrast, the male proximal tubules reabsorbed only 53% of the filtered Na^+^; the female proximal tubules even less, only 34%. Consequently, downstream segments in mice must handle a larger fraction of the filtered Na^+^. In particular, the thick ascending limbs mediate 44% of the filtered Na^+^ in male mouse and 64% in female, significantly more than the 23% in male rat and 37% in female rat. Since Na^+^ transport drives the reabsorption of Cl^-^ and water, and other electrolytes similar interspecies difference is also seen in the segmental transport of water and other solutes; see Fig. 2.

Further comparison can be found in the fractional delivery and excretion rates of water, Na^+^, and K^+^, which are summarized in Table 2. Predictions for the rat models have been previously validated (9, 16). Predictions for the male mouse model are consistent with micropuncture data (20, 34, 39).

**Table 2.** Predicted fraction deliveries and excretions (%) in the rat and mouse nephron models.

|  | Male rat | Female rat | Male mouse | Female mouse |
| --- | --- | --- | --- | --- |
| **Late proximal tubule** |  |  |  |  |
| **Volume** | 21 | 36 | 39 | 54 |
| **Na<sup>+</sup>** | 27 | 43 | 47 | 66 |
| **K<sup>+</sup>** | 29 | 47 | 51 | 84 |
| **Late distal convoluted tubule** |  |  |  |  |
| **Volume** | 13 | 17 | 19 | 25 |
| **Na<sup>+</sup>** | 42 | 26 | 40 | 36 |
| **K<sup>+</sup>** | 15 | 20 | 20 | 29 |
| **Urine** |  |  |  |  |
| **Volume** | 1.1 | 1.4 | 1.1 | 1.3 |
| **Na<sup>+</sup>** | 1.0 | 1.3 | 0.9 | 1.1 |
| **K<sup>+</sup>** | 25 | 27 | 26 | 36 |

### Males versus females

The sexual dimorphism in renal transporters in rats and mice exhibits similarities as well as differences. In both species, the abundance of key transporters in the proximal tubules is lower in females. Notably, there is are fewer NHE3 overall in female mice, and more phosphorylated (inactive) NHE3 in female rat, both of which implies lower NHE3 activity in females. The abundance of claudin-2 is also lower both females. One notable difference between the two species is the sex differences in proximal tubule AQP1 expression, which is 40% higher in female mice compared to males, but 60% lower in female rats. These differences in proximal transporter abundance are represented in the model; see Table 1. The lower NHE3 activity in females appear to have the most dominating effect: the models predict that a larger fraction of the filtered Na^+^ is reabsorbed in the male species relative to females, as noted above. Because water transport is driven by Na^+^ reabsorption, even though the female mouse expression more AQP1, a significantly larger fraction of the filtered volume is reabsorbed along the proximal tubules in the male mouse (61%) compared to female (46%).

The resulting proportionally higher Na^+^ load and volume in females is handled by the distal segments, which express a higher abundance of key transporters, including more NKCC, NCC, and claudin-7 in mice, and more ENaC, NCC, ENaC, and claudin-7 in rats. These sex differences are represented by setting appropriate transport parameter values in the models; see Table 1.

### Responses to Na^+^ load

Veiras et al. (37) reported that the response to an acute Na^+^ load differs between rats and mice, and between the two sexes. Specifically, given the same saline bolus volume relative to body weight, diuretic and natriuretic responses were more rapid in female rats, with the fractional excretion of the bolus more than twice in female rats compared to males at 3 h. In contrast, no significant difference was found between the excretion rates of male and female mice. We conducted simulations to determine whether the baseline model transport parameters would predict a similar sex difference in rats, and no difference in mice. To simulate a saline load, we increase SNGFR by 50% in the male rat and mouse models (2, 4), and by 65% in the female rat and mouse models, to take into account the greater salt-loading induced increases in macula densa NOS1β expression and activity in females than in males (42). All other parameters remained unchanged.

The model predicts some degree of glomerulotubular balance in all four models, with proximal tubular Na^+^ transport increases in response to the higher Na^+^ load (Fig. 3). Water and Cl^-^ transport primarily follows Na^+^; the model predicts significant increases in proximal tubular water and Cl^-^ reabsorption in all models (Fig. 3). Despite the enhanced proximal reabsorption, Na^+^ delivery to the thick ascending limb is about twice the baseline values. Consequently, Na^+^ reabsorption along the thick ascending limb is also predicted to increase. Na^+^ load is predicted to substantially impact K^+^ transport along the thick ascending limb and connecting tubule. K^+^ reabsorption approximately doubles along the thick ascending limb, whereas K^+^ is secreted along the connecting tubule, and that secretion is predicted to increase markedly (Fig. 3).

**Fig. 3.**
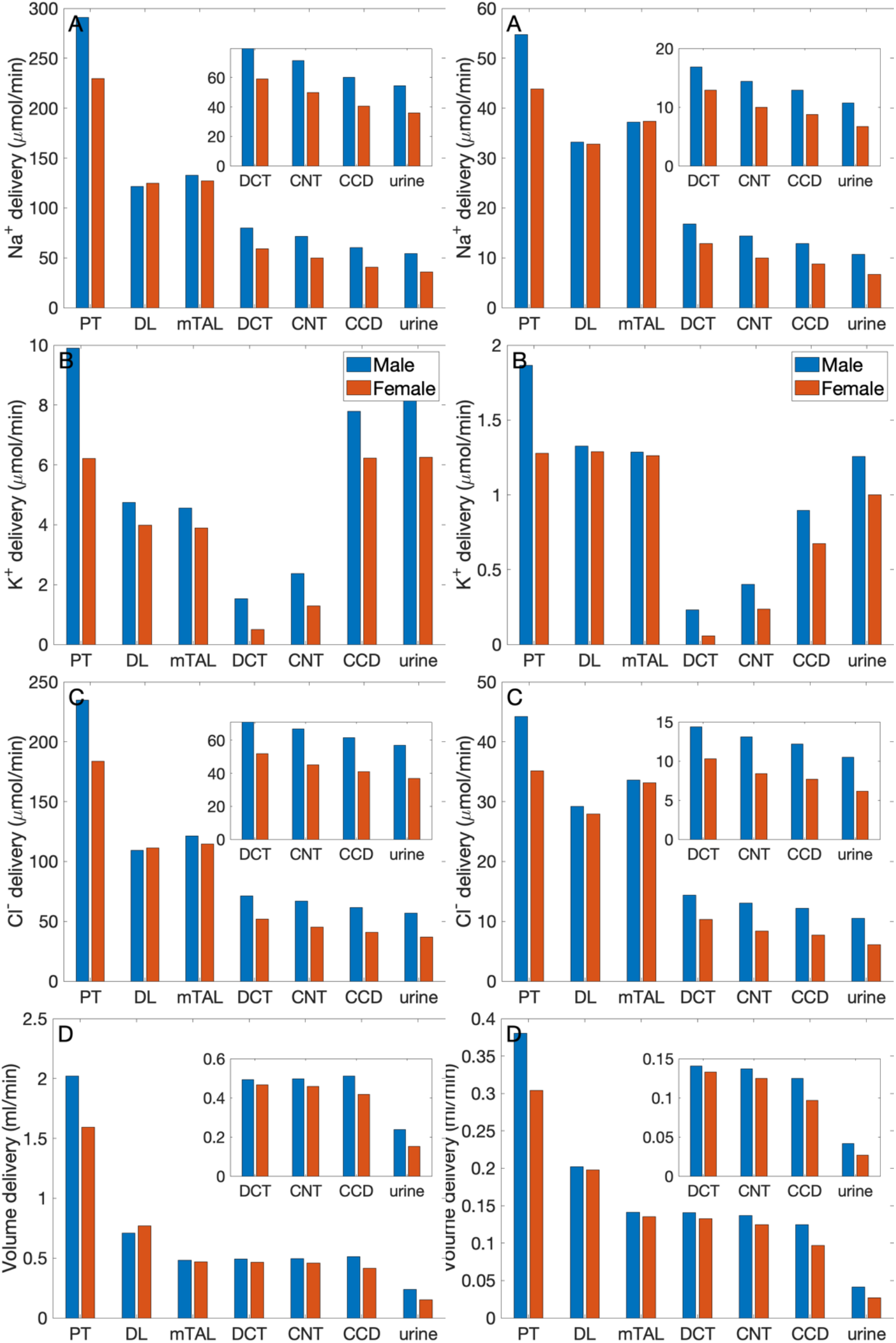
Delivery of key solutes and fluid to the beginning of individual nephron segments in male and female rats (left column) and mice (right columns) following a saline load. Notations are analogous to Fig. 2. The predicted excretion rates are lower for the female models relative to the respective male model. However, when scaled by bolus volume, the female rat model excretes the bolus faster than the male rat model.

Taken together, the above transport processes result in urine flow of 241 and 194 *μ*L/min, respectively, in male and female rats, and 41 and 38 *μ*L/min, respectively, in male and female mice, given per kidney. Urinary Na^+^ excretion is predicted to be 54 and 47 *μ*mol/min, respectively, for each male and female rat kidney, and 11 and 10 *μ*mol/min, respectively, for each male and female mouse kidney. Assuming that a female rat weighs about one-half that of the male rat (at 10−12 wk, Envigo.com), then the saline bolus volume simulated in the female model is one-half that of the male model. Thus, the female-to-male ratio of Na^+^ excretion (denoted *E*_Na_) as a fraction of saline load is given by (female *E*_Na_/male *E*_Na_)/(female saline volume/male saline volume) = (47/54)/0.5 = 1.74. A mature male C56BL/6 mouse weighs approximately 30% more than an age-matched female (16 weeks; the Jackson Laboratory). The analogous ratio for mice is given by (10/11)×1.3 = 1.18. Thus, the model predicts an interesting inter-species difference: that female rats excrete a saline load significantly faster than male rats, whereas male and female mice had more similar responses. That result is consistent with findings by Veiras et al. (37) Similarly, the female-to-male ratios of fractional volume excretion are predicted to be significantly higher for rats (1.58) than for mice (1.20).

### Segmental contributions to interspecies differences in nephron transport

As previously noted, between the rat and the mouse, there are notable differences in the fractional solute and water transport by individual nephron segments. These differences are found in both males and females, and are a result of the differences in transporter patterns of the two species. To understand the functional implications of these inter-species differences, we first conducted simulations to study how differences in transporter patterns of individual segments affect solute and water transport along those segments, as well as urinary excretion. Specifically, we set the transporter and channel activities of individual nephron segments of the mouse model to the corresponding rat values (of the same sex), and computed the resulting changes in segmental transport and urinary excretion. Six “segments” were considered: (i) proximal convoluted and straight tubules, (ii) medullary and cortical thick ascending limbs, (iii) distal convoluted tubule, (iv) connecting tubule, (v) cortical and outer-medullary collecting duct, and (vi) inner-medullary collecting duct. Only transporter and channel activities were modified, but not other parameters such as SNGFR and segmental lengths and diameters. For example, for the proximal tubule simulations, NHE3 activity along the proximal convoluted and straight tubules in the mouse models was increased by 47% to agree with the rat values (to eliminate the 32% difference between rat and mouse; see Table 1). That would be the only change.

Figure 4 shows percentage changes in Na^+^, K^+^, and water transport along each mouse nephron segment if its transporter abundance were the same as a sex-matched rat. These results answer the question: *How do these inter-species differences affect transport along that particular segment*? However, due to the large differences in fractional transport among segments (e.g., under baseline conditions, the proximal tubules of the male mouse reabsorb almost 20 times more Na^+^ transport the cortical and outer-medullary collecting ducts), these percentage changes do not translate directly into absolute changes, and cannot be compared between segments. Also, transport along downstream segments was also affected (results not shown). To assess impacts on overall kidney function, resulting changes in Na^+^ excretion, K^+^ excretion, and urine volume are also included in Fig. 4. These results answer the question: *How do mouse-specific transporter activity values along individual segments impact overall transport*?

**Figure 4.**
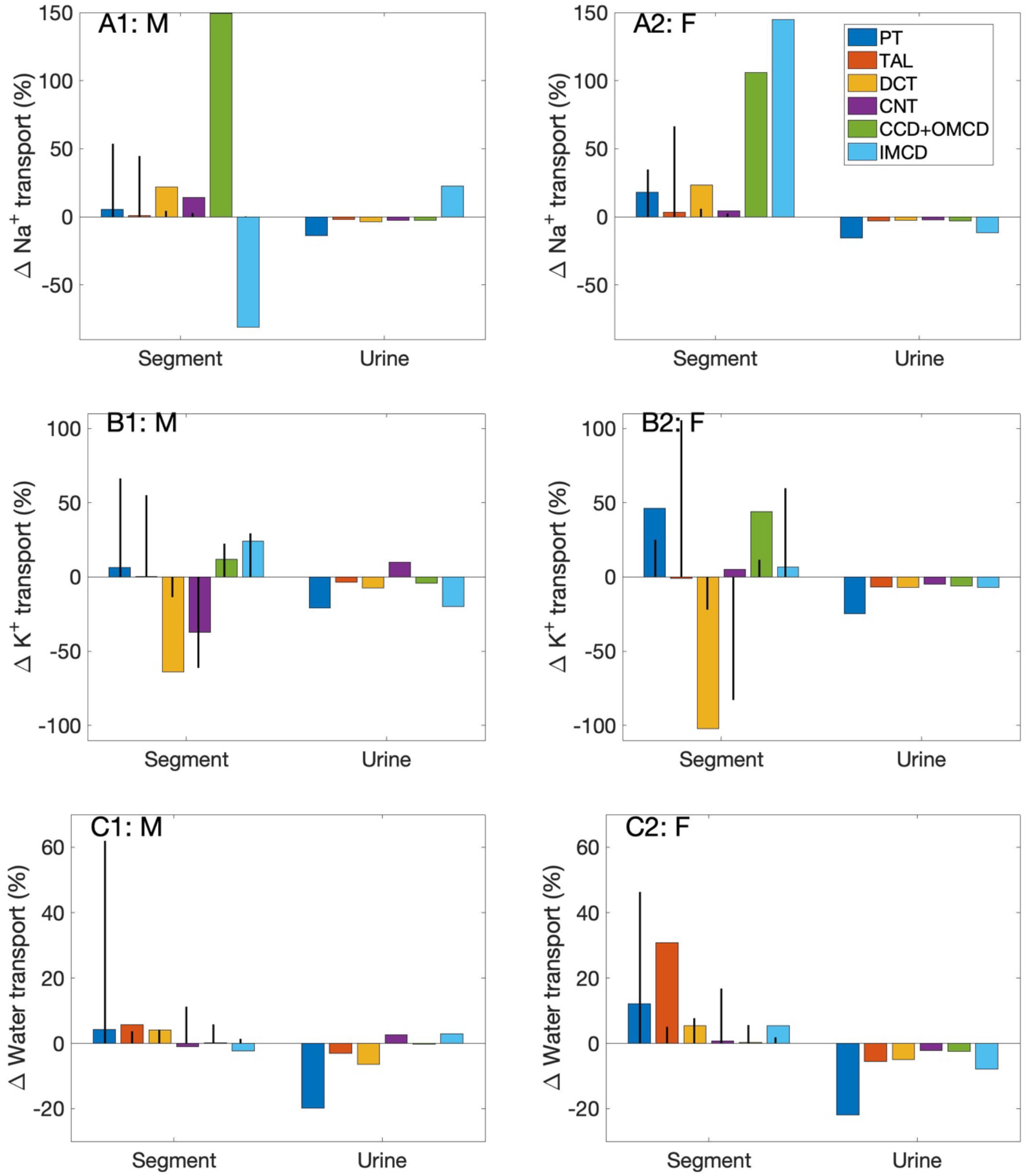
Effects on segmental transport and urine excretion if transporter abundance in individual segments was set to values in the sex-matched rat. Results show (i) fractional changes in in Na^+^ transport (A1, A2), K^+^ transport (B1, B2), and water transport (C1, C2) along each nephron segment in the male mouse (left column) and female mouse (right column) if transporter abundance in individual segments were set to values in the sex-matched rat, and (ii) Na^+^ excretion, K^+^ excretion, and urine output. Six segments were considered individually: proximal tubule (PT), thick ascending limb (TAL), distal convoluted tubule (DCT), connecting tubule (CNT), cortical and outer-medullary collecting ducts (CCD+OMCD), and inner-medullary collecting duct (IMCD). Middle bars indicate fractional transport mediated by that segment under baseline conditions.

Changes in the proximal tubules and inner-medullary collecting ducts have the largest effect on urinary excretions and output. In the mouse proximal tubule, NHE3 is downregulated compared to the rat. If a mouse proximal tubule exhibits the same NHE3 activity as a sex-matched rat, proximal tubule transport would significantly elevate and, despite some compensation along downstream segments, Na^+^ excretion, K^+^ excretion, and urine output would all decrease by up to 20%. Changes in the proximal tubules have a large effect on nephron function because, except in the female mouse (more below), the proximal tubules via the NHE3 mediate the majority of Na^+^ transport. In the female mouse, most of the filtered Na^+^ is reabsorbed by the thick ascending limbs. However, the difference in thick ascending limb transporter abundance between the rat and mouse is small, only a 13% reduction in Na^+^-K^+^-ATPase activity. When Na^+^-K^+^-ATPase activity along the thick ascending limbs is increased to rat level, the models predict only minor reductions in Na^+^ excretion, K^+^ excretion, and urine output.

Changes in the inner-medullary collecting duct also have a large effect on nephron function even though that segment is responsible for only a minute fraction of overall nephron transport because a number of transporter parameters are assumed to be different between the rat and the mouse (NHE3, Na^+^-K^+^-ATPase, and H^+^-K^+^-ATPase activity and paracellular Na^+^ permeability), and, as importantly, it is the last segment with no downstream segments to compensate. These changes have competing effects on electrolyte and water transport. Thus, the net effects on urinary excretions and output depends on the solute and sex. See Fig. 4.

### A closer look at sex differences in the rat versus mouse nephron

Sexual dimorphism in transport pattern is qualitatively similar between the rat and mouse, except for a few transporters. Based on the immunoblotting results by Verias et al. (37), the present models assume (i) proximal tubule apical water permeability (P_f_) is lower in female rat compared to male rat, but higher in female mouse compared to male mouse; (ii) Na^+^-K^+^-ATPase activity is doubled in female rat from males along the thick ascending limbs, distal convoluted tubules, and connecting tubules, but the analogous sex difference is much smaller in mice; (iii) ENaC activity is assumed to be significantly higher in female rat relative to male, but there is no sex difference in mice. See Table 1. *What are the function implications of these interspecies differences in sex differences?* To answer that question, we conducted three simulations, in which selective sex differences in the mouse models were made rat-like. Specifically, (i) proximal tubule P_f_ in the female mouse was set to 36% lower than male mouse like the rats (instead of 40% higher); (ii) Na^+^-K^+^-ATPase activity along the thick ascending limbs, distal convoluted tubules, and connecting tubules in female mouse is set to twice the male mouse values (instead of small to no change); (iii) ENaC activity along the connecting tubules, cortical and outer-medullary collecting ducts of the female mouse is set to 30%, 50%, and 20%, respectively, above male values (instead of no difference).

As expected, all these changes increased Na^+^ reabsorption along the affected segments; see Fig. 5. Water transport increased as well. Note that while the percentage increase in water transport for the “thick ascending limb Na^+^-K^+^-ATPase activity” case is nearly 30%, the net increase is small because the thick ascending limbs are nearly water impermeable. K^+^ transport increased as well, except when thick ascending limb Na^+^-K^+^-ATPase activity was increased, which limits K^+^ reabsorption along the thick ascending limbs. All changes suppressed urinary Na^+^ and K^+^ excretion and urine output, except when ENaC activity is increased. That modification increases K^+^ secretion and excretion.

**Figure 5.**
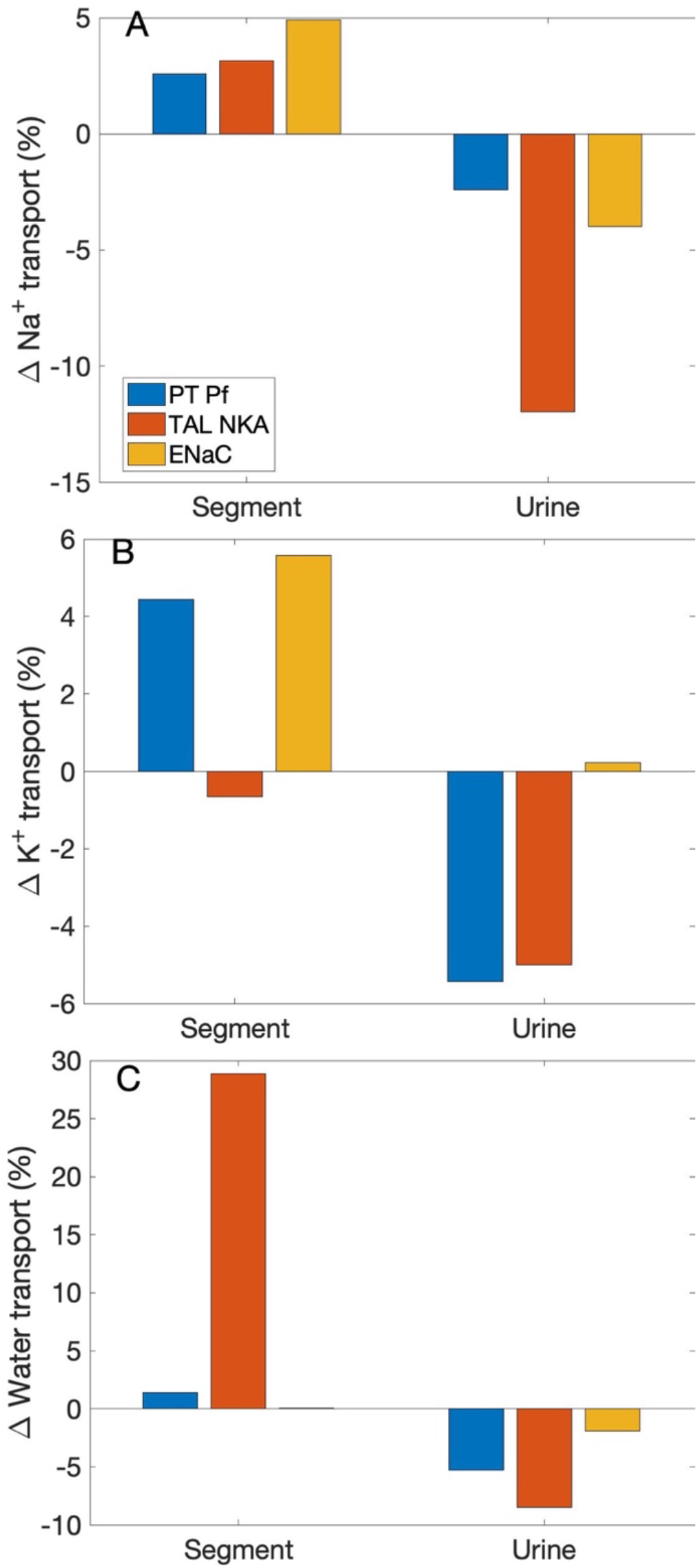
Effects on segmental transport and urinary excretion in the female mouse if selected sex differences in mice were change to corresponding differences in the rat. Three sex differences that are most qualitatively different between the two species were considered: (i) proximal tubule apical water permeability (“PT Pf”), (ii) thick ascending limb Na^+^-K^+^-ATPase (“TAL NKA”), and (iii) ENaC. Results show fractional changes in Na^+^ transport (A), K^+^ transport (B), and water transport (C) along affected segments, as well as the corresponding changes in urinary excretion rates and volume.

## Discussion

Physiology is fascinating in its many layers of complexity. Even within a species, one finds that the structures and functions of organ and tissue systems may be affected by a multitude of factors, including sex and gender, age, time of day or year, and race or strain. The present study analyzes kidney function along two dimensions, sex and species. To accomplish that goal, we developed and applied sex-specific computational models for nephron transport in the rat and mouse kidney. The models represent the sexual dimorphism in renal morphology, hemodynamics, and transporter abundance (23, 36). Model simulations were conducted to compare kidney function between the rat and mouse, between males and females.

Before discussing the physiological insights gleaned from the simulation results, we note that a notable contribution of this study is the development of epithelial transport models of the mouse kidney. The mouse is the animal model used in over 95% of animal studies, given its small size and the similarity it shares with human’s genome (99%). Genetically modified mouse models have been essential for understanding physiological mechanisms, as well as the pathogenesis, progression, and treatments of diseases. Genetically manipulated animals almost always develop compensatory response, which can muddle the results. To help distinguish response to the intended genetic manipulation from compensation, computational models can used to simulate “clean” genetic modification experiments. However, while most of the genetical modification animal models are mice, computational models of the mammalian kidney have been focused on the rat, because of the more abundant morphological, electrochemical, and micropuncture data. Recently, analogous computational models were developed for humans, for their higher and more relevant clinically value, despite the sparsity of measurements needed to determine model parameters. In contrast, no such models exist, until now, for the mouse. In the absence of a mouse computational model, theoretical analysis was often performed using a rat model to explain experimental observations made in mouse studies (e.g., (15, 17, 38)). That is, clearly, less than ideal, given the plentiful inter-species differences in morphology, hemodynamics, and molecular structure, some of which have been discussed in this study. The mouse nephron models developed in this study finally allows us to supplement genetically modified mouse kidney experiments with *in silico* studies using computational models of the mouse kidney. Recognizing that sex is a key biological variable, the mouse nephrons models, like their rat counterparts, are sex specific. Thus, not only do these mouse models yield relevant and accurate predictions for both sexes, but they can also be used to extrapolate experimental findings obtained in one sex to the other.

Another goal of this study is to understand the physiological implications of the differences in renal transporter distribution between female and male nephrons. Both the rat and mouse nephrons exhibit lower Na^+^ and water transporters and lower fractional reabsorption in the proximal nephron of females versus males, coupled with more abundant transporters in renal tubule segments downstream of the macula densa (36). Fine control of Na^+^ excretion, important in the regulation of body electrolyte, volume, and blood pressure, is accomplished by the tight regulation of Na^+^ transport along the distal tubular segments (7). Hence, the larger contribution of the distal nephron segments to total renal Na^+^ transport in females may better equip them for the fluid and electrolyte retention and adaptations demanded of them during pregnancy and lactation. Results of our recent modeling studies (30) suggest that the female renal transporter pattern, together with the pregnancy-induced adaptations, make possible the fluid, Na^+^, and K^+^ retention required for a healthy pregnancy. Those models were formulated for a rat in mid- and late-pregnancy. Extending the model to a pregnant mouse, when the necessary transporter expression data become available, might shed light into whether the female mouse transporter abundance pattern, despite its differences from that of the female rat, would also meet the enhanced fluid and electrolyte demands in pregnancy.

Notwithstanding the similarities in the sex differences in transporter abundance patterns in the rat and mouse nephrons, there are notable differences in the proximal tubules (AQP1) and distal segments (NKCC2, Na^+^-K^+^-ATPase, and ENaC). These inter-species differences may provide a partial explanation for the two rodents’ different response to an acute Na^+^ load. Veiras et al. (36) reported that female rats excrete an acute Na^+^ load faster than male rats. Specifically, given the same saline bolus volume relative to body weight, diuretic and natriuretic responses were more rapid in female rats, with female rats excreting ∼30% and male rats <15% of the saline bolus at 3 h. In contrast, no significant difference was observed between the male and female mice. That difference in the rat versus mouse response is likely not fully explained by the discrepancies in their baseline kidney structure and function, as an acute saline load will likely trigger alternations in renal transporter expression, which have not been measured separately for each sex and are thus not incorporated in the present simulations. Furthermore, extrarenal systems that participate in the pressure-natriuresis response, such as the renin-angiotensin system, exhibit differences between the two species and between sexes. Thus, a full understanding of each species’ sex-specific response to a saline load and other hypertensive stimuli requires a more comprehensive computational model (e.g., (1, 18)). Given that the kidney maintains the balance of not only Na^+^ but other electrolytes and acid-base, it would be interesting to better characterize any sex dimorphism in the two species’ response to K^+^ load or other stimuli (e.g., (8)) differs significantly, and if that can be explained by differences in their transporter abundance patterns.

It is our hope that our computational kidney models can be a useful tool in synthesizing and analyzing the large amount of data emerging from different experimental studies. The ultimate goal is to utilize in silico studies to advance our understanding of kidney physiology and pathophysiology, and support therapeutic drug development. With that goal in mind, limitations of the present models must be acknowledged. As noted above, kidney function is modulated not only by sex but many other factors as well. For instance, numerous physiological functions exhibit circadian oscillations (25), and the kidney is no exception. Renal plasma flow, GFR, and tubular reabsorption and/or secretion processes have been shown to peak during the active phase and decline during the inactive phase (5). Interestingly, but perhaps not surprisingly, the circadian regulation of renal transporter expression level exhibits sex difference (6). Incorporating circadian regulation of filtration and transport into the computational models, expanding upon what was done for an isolated proximal tubule cell model (38), would make for more chronologically accurate predictions of kidney function, and provide a new tool for chronopharmacological studies.

